# A powerful approach to estimating annotation-stratified genetic covariance using GWAS summary statistics

**DOI:** 10.1101/114561

**Authors:** Qiongshi Lu, Boyang Li, Derek Ou, Margret Erlendsdottir, Ryan L. Powles, Tony Jiang, Yiming Hu, David Chang, Chentian Jin, Wei Dai, Qidu He, Zefeng Liu, Shubhabrata Mukherjee, Paul K. Crane, Hongyu Zhao

## Abstract

Despite the success of large-scale genome-wide association studies (GWASs) on complex traits, our understanding of their genetic architecture is far from complete. Jointly modeling multiple traits’ genetic profiles has provided insights into the shared genetic basis of many complex traits. However, large-scale inference sets a high bar for both statistical power and biological interpretability. Here we introduce a principled framework to estimate annotation-stratified genetic covariance between traits using GWAS summary statistics. Through theoretical and numerical analyses we demonstrate that our method provides accurate covariance estimates, thus enabling researchers to dissect both the shared and distinct genetic architecture across traits to better understand their etiologies. Among 50 complex traits with publicly accessible GWAS summary statistics (N_total_ ≈ 4.5 million), we identified more than 170 pairs with statistically significant genetic covariance. In particular, we found strong genetic covariance between late-onset Alzheimer’s disease (LOAD) and amyotrophic lateral sclerosis (ALS), two major neurodegenerative diseases, in single-nucleotide polymorphisms (SNPs) with high minor allele frequencies and in SNPs located in the predicted functional genome. Joint analysis of LOAD, ALS, and other traits highlights LOAD’s correlation with cognitive traits and hints at an autoimmune component for ALS.

## Introduction

Genome-wide association study (GWAS) has been a success in the past 12 years. Despite a simple study design, GWAS has identified tens of thousands of robust associations for a variety of human complex diseases and traits. Based on the GWAS paradigm, linear mixed models, in conjunction with the restricted maximum likelihood (REML) algorithm, have provided great insights into the polygenic genetic architecture of complex traits [1-3]. The cross-trait extension of linear mixed model has further revealed the shared etiology of many different traits [4]. Compared to traditional, family-based approaches, these methods do not require all the traits to be measured on the same cohort, and therefore make it possible to study a spectrum of human complex traits using independent samples from existing GWASs [5, 6]. Recently, Bulik-Sullivan et al. developed cross-trait LDSC, a computationally efficient method that utilizes GWAS summary statistics to estimate genetic correlation between complex traits [7]. LDSC is a major advance. As summary statistics from consortium-based GWASs become increasingly accessible [8], it provides great opportunities for systematically documenting the shared genetic basis of a large number of diseases and traits [9, 10]. However, large-scale inference sets a high bar for both estimation accuracy and statistical power. Furthermore, existing methods do not allow explicit modeling of functional genome annotations. As shown in later sections, the estimated genetic correlations in many cases are neither statistically significant nor easy to interpret.

To address these challenges, there is a pressing need for a statistical framework that provides more accurate covariance and correlation estimates and allows integration of biologically meaningful functional genome annotations. The method of moments has recently been shown to outperform LDSC in single-trait heritability estimation [11]. Integrative analysis of GWAS summary statistics and context-specific functional annotations has provided novel insights into complex disease etiology through a variety of applications [12-14]. In this paper, we introduce GNOVA (GeNetic cOVariance Analyzer), a principled framework to estimate annotation-stratified genetic covariance using GWAS summary statistics. Through extensive numerical simulations, integrative analysis of 50 complex traits, and an in-depth case study on late-onset Alzheimer’s disease (LOAD [MIM: 104300]) and amyotrophic lateral sclerosis (ALS [MIM: 105400]), we demonstrate that GNOVA provides accurate covariance estimates and powerful statistical inference that are robust to linkage disequilibrium (LD) and sample overlap. Furthermore, we show that annotation-stratified analysis enhances the interpretability of genetic covariance and provides novel insights into the shared genetic basis of complex traits.

## Material and methods

### Statistical model

Here we outline the genetic covariance estimation framework. The complete derivation, detailed justification for all approximations, and theoretical proofs are presented in the **Supplementary Notes**. In short, the genetic covariance that we aim to estimate is the covariance between the genetic effects of a group of single nucleotide polymorphisms (SNPs) on two complex traits. When functional genome annotations are present, we allow such covariance to vary in different annotation categories. Specifically, we define K functional annotations *S*_1_, *S*_2_, …, *S*_*K*_(e.g. protein-coding genes and non-coding regions), whose union covers the entire genome; assume two studies share the same list of *m* SNPs; and assume two standardized traits *y*_1_ and *y*_2_ follow the linear models below:

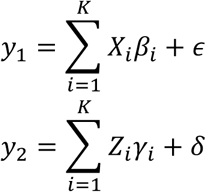

where *X*_*i*_ and *Z*_*i*_ denote the standardized genotype matrices defined through annotation *S*_*i*_. Random effects terms *β*_*i*_ and *γ*_*i*_ denote the corresponding genetic effects for each annotation category. SNPs’ genetic effects on two traits follow an annotation-dependent covariance structure:

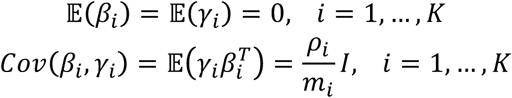

where *m*_*i*_ and *ρ*_*i*_ denote the total number of SNPs and the total genetic covariance in annotation category *S*_*i*_, respectively. Random variables *ε* and *δ* denote the non-genetic effects. Of note, this notation implicitly assumes the genetic covariance to follow an additive structure in regions where functional annotations overlap.

In practice, two different GWASs often share a subset of samples. Without loss of generality, we assume *N*_1_ and *N*_2_ to be the sample sizes of two studies and the first *N*_*s*_ samples in each study are shared. To account for the non-genetic correlation introduced by sample overlapping, we allow random error terms *ε* and *δ* to be correlated:

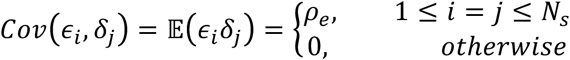

We note that our model does not require any additional assumption on the heritability structure of either trait.

### Estimation of covariance parameters using the method of moments

To estimate genetic covariance parameters (i.e. *ρ*_*i*_, *i* = 1, …, *K*), we developed an analysis framework based on the method of moments. First, we derive equations that relate the population moments to the parameters of interest. For an arbitrary *N*_1_×*N*_2_ matrix *A*, we study the expectation of 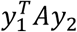. It can be shown that

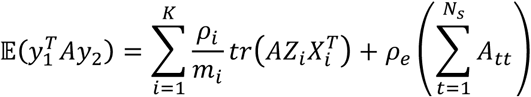

Here, quantity *A*_*tt*_ denotes the t^th^ diagonal element of matrix *A*. Since there are K+1 parameters in total in the model (K genetic covariance parameters and *ρ*_*e*_), we build a linear system of K+1 equations by plugging in K+1 different matrices *A*_1_,…, *A*_*K* + 1_ into the equation above. Further, we approximate 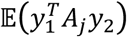 using the sample moments, i.e. the observed value 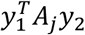, and get the following equations.

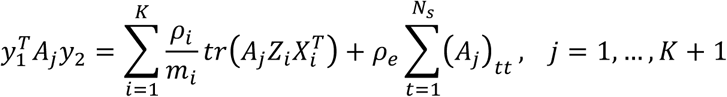

Solving this linear system of K+1 equations would get us the method of moments estimators for genetic covariance.

### Choices of matrix *A*

The method of moments estimation procedure described above works for arbitrary matrices. However, it is critical and non-trivial to choose A in practice. Since individual-level genotype and phenotype data from consortium-based GWASs are in many cases difficult to access, it is of practical interest to estimate genetic covariance based on summary statistics only. To achieve this goal, we define the first K matrices as:

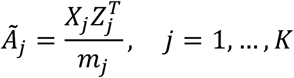

Plugging in these matrices, the first K equations become:

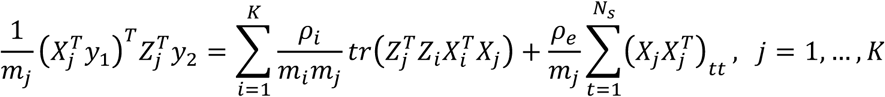

The equality is based on the property of trace and the fact that first *N*_*S*_ samples are shared between two studies. These equations can be further approximated by (**Supplementary Notes**):

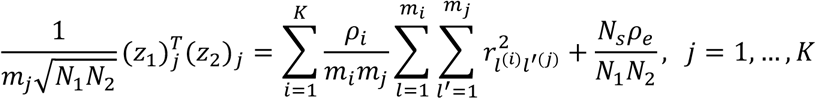

Here, 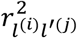 denotes the LD between the *l* ^th^ SNP from category *S*_*i*_ and the (*l*′)^th^ SNP from category *S*_*j*_; *z*_1_ and *z*_2_ denote the z-scores of SNP-level associations from two GWASs; (*z*_1_)_*j*_ and (*z*_2_)_*j*_ represent subsets of z-scores corresponding to the SNPs in annotation category *S*_B_. LD can be estimated using an external reference panel. However, if samples in two studies have different ancestries, 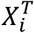*X*_*j*_ and 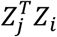 need to be estimated separately using two reference panels. When such reference panels do not exist, individual-level genotype data for a subset of study samples may be needed.

Next, we study the (K+1)^th^ equation. We define:

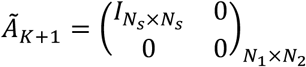

Divide *N*_1_*N*_2_ on both sides of the (K+1)^th^ equation, and we get:

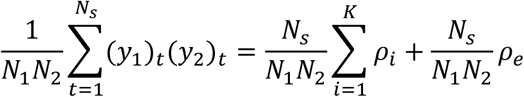

Since *ρ*_1_, …, *ρ*_%_ are the parameters of interest, we subtract the (K+1)^th^ equation from the first K equations, and remove *ρ*_*K*+1_ from the linear system. We denote the remaining K equations in matrix form:

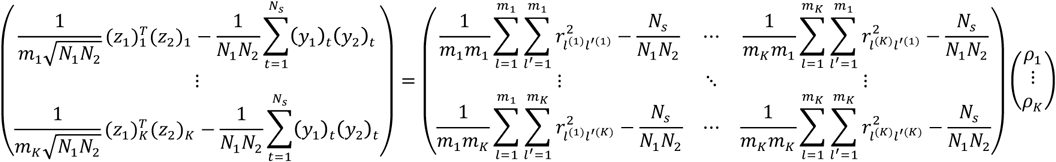

When the sample sizes of both GWASs are large and the sample overlap between two studies is moderate, the K equations can be approximated by:

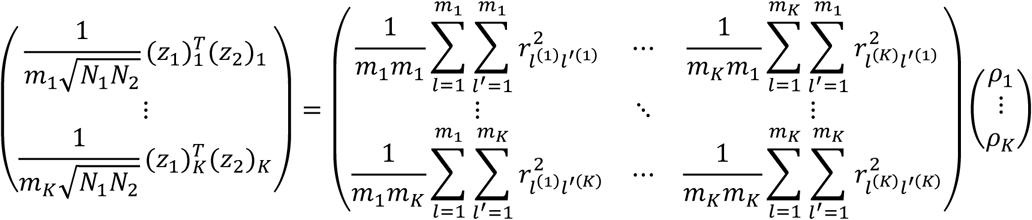

We define

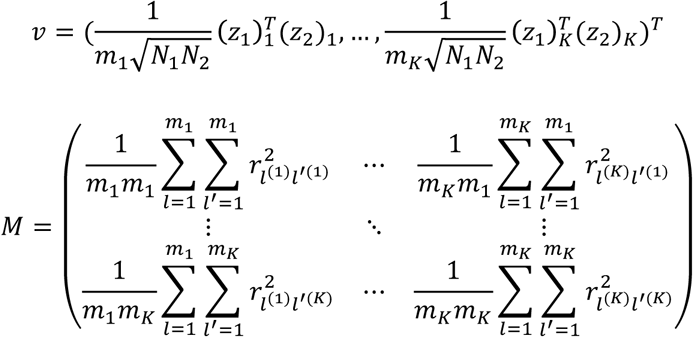

Then, the point estimate of covariance parameters can be denoted as

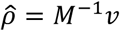

Importantly, *M* can be estimated using a reference panel (e.g. 1000 Genomes Project [15]) and *v* is only based on GWAS summary statistics. Of note, the same estimation framework can be directly applied to ascertained case-control studies as well (**Supplementary Notes**).

### Special cases

#### 1) Two independent GWASs

If samples from two GWASs do not overlap, then the non-genetic effects *ε* and *δ* are independent and only K equations are needed for estimating covariance estimators. We still define 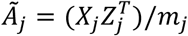 for j=1,…,K. That gives us the same covariance estimator:

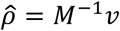

#### 2) No annotation stratification

If no functional annotation is present, it can be shown that

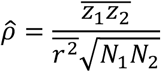

Here, 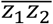 is the average product of z-scores from two GWASs; 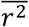 is the average LD across all SNP pairs in the study. Under the non-stratified scenario, this estimator can be seen as a two-trait extension of the heritability estimator proposed in [16].

#### 3) Two GWASs with substantial sample overlap

If the two GWASs have substantial sample overlap, some approximations we have applied in previous sections would fail (**Supplementary Notes**). The problem gets down to solving the following equations.

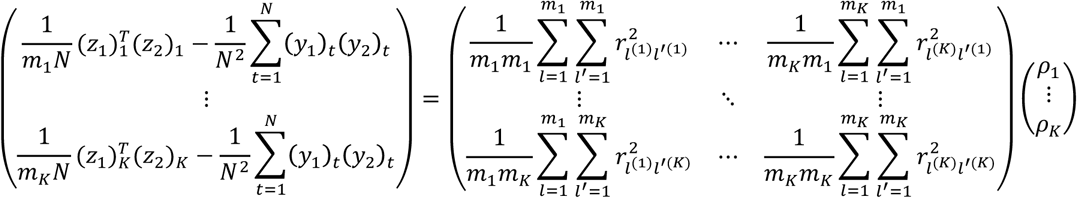

Therefore,

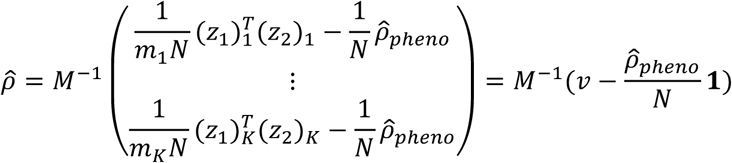

where the phenotypic correlation 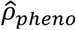 can be either acquired from the literature or estimated using computational methods [7, 17, 18] (**Supplementary Notes**).

### Remarks on overlapping functional annotations

When functional annotations overlap, the covariance parameter *ρ* is not the real quantity of interest. Instead, the total covariance in each annotation category is more biologically meaningful and can be estimated using the weighted estimator

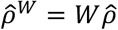

where w is a *K*×*K* matrix with element

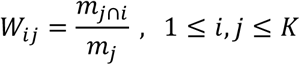

Here, *m*_*j*∩*i*_ denotes the number of SNPs in region *S*_*i*_ ∩ *S*_B_.

### Theoretical properties

In this section, we establish the statistical optimality of our estimator by showing that it is “almost” the unbiased estimator with minimum variance. Here we state all the propositions. See **Supplementary Notes** for detailed proofs. Assume *y*_1_ and *y*_2_ follow a multivariate normal distribution:

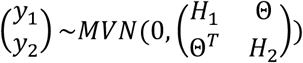

We begin with calculating the variance of the quadratic form-like quantity 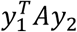.

#### Proposition 1.

Let *A* be an *N*_1_×*N*_2_ matrix. Then 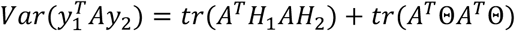.

It can be shown that the second part, i.e. *tr*(*A*^T^Θ*A*^T^Θ, is very small compared to the first term *trA*^T^*H*_1_*AH*_2_ in real GWAS data (**Supplementary Notes**).

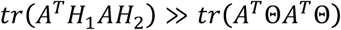

With this in mind, the following claim is approximately true.

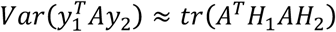

Next, we define a matrix *A*∗, and show that *A*∗ minimizes *tr*(*A*^1^ *H*_1_ *AH*_2_) under some conditions. Based on the argument above, *A*∗ “almost” minimizes 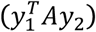 too.

#### Proposition 2.

Assume two GWASs do not share samples. We define the following quantities.

i. Let *p*=(*p*_*1*_,…*p*_*k*_)^*T*^ be an arbitrarily given K-dimensional vector;
ii. Let *S* be a *K*×*K* symmetric matrix with element 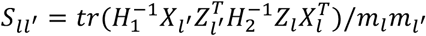 for 1 ≤ *l*, *l*′ ≤ *K*;
iii. Let *λ*=(*λ*_*1*_,… *λ*_*k*_)^*T*^ be a vector such that *S* λ = *p*;
iv. Define 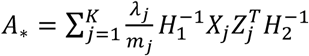.

Then, we have:

1. 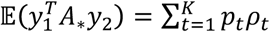
2. Let *A* be a matrix such that 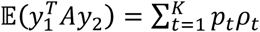. Then, 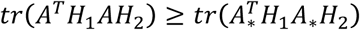.

Proposition 2 tells us that given arbitrary *p*=(*p*_*1*_,…*p*_*k*_)^*T*^ if ∃ *λ*=(*λ*_*1*_,… *λ*_*k*_)^*T*^ such that *S*λ = *p*, then 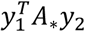 is an unbiased estimator for 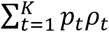. Furthermore, among all unbiased estimators with the form 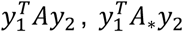 has the minimum value of 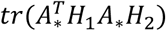, hence “almost” the minimum variance 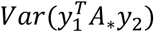. Interestingly, by carefully choosing *p* and *λ*, we can let *A*∗ equal the Ã matrix we have been using throughout the paper. Therefore, we have the following corollary.

***Corollary 1.*** We assume:

i. Two GWASs do not overlap;
ii. The samples in each study are completely independent;
iii. True LD in both studies (i.e. *Z*^*T*^*Z* and *X*^*T*^*X*) is known.

Consider all matrices *A* that suffice

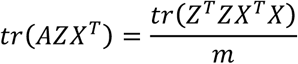

We define

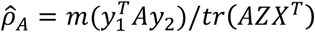

Then, 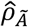 with Ã = (*XZ*^*T*^)/*m* has the lowest variance.

Similarly, we could extend these results to annotation-stratified scenarios (**Supplementary Notes**). These results show that although we initially defined *Ã*_*j*_ for the purpose of simplifying calculation, the derived covariance estimator actually enjoys some good theoretical properties.

### Variance estimation using block-wise jackknife

Following [7], we apply a block-wise jackknife approach to estimate the variance. We divide the genome into *b* (e.g. *b* = 200) blocks *B*_1_, …, *B*. Let

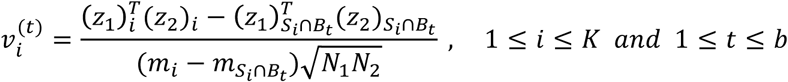

Here, subscript *S*_*i*_ ∩ *B*_*t*_ indicates the subset of SNPs in both functional annotation *S*_*i*_ and block *B*_*t*_. Then, *Cov*(*v*) is estimated as:

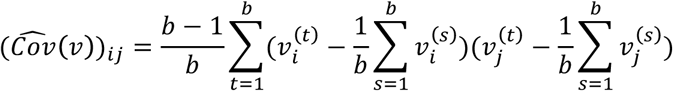

Therefore, we get

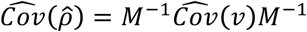

If annotations overlap,

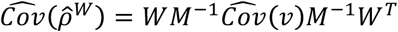

Finally, the test statistic for each covariance parameter is

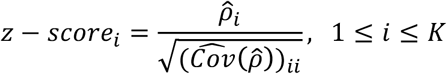

When annotations overlap,

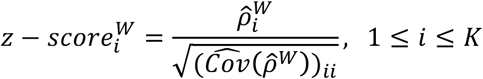

### Genetic correlation

We provide genetic correlation estimates for non-stratified analysis.

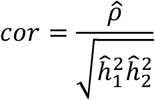

We use the estimator proposed in [16] to estimate heritability for each trait.

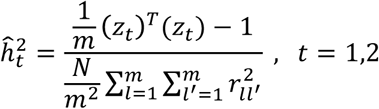

When functional annotations are present, the true heritability in each annotation category may be small. Although methods for estimating annotation-stratified heritability have been proposed [11, 12], they may provide unstable, sometimes even negative heritability estimates, especially when a number of annotation categories are related to the repressed genome. When true heritability is low, variability in the denominator will have great impact on genetic correlation estimates. Therefore, we use genetic covariance as a more robust metric when performing annotation-stratified analysis.

### Simulation settings

We simulated quantitative traits using real genotype data from the WTCCC1 cohort. We removed individuals with genetic relatedness coefficient greater than 0.05 and filtered SNPs with missing rate above 1% and/or MAF lower than 5% in samples with European ancestry from the 1000 Genomes Project [15]. In addition, we removed all the strand-ambiguous SNPs. After quality control, 15,918 samples and 254,221 SNPs remained in the dataset. Each simulation setting was repeated 100 times.

#### Setting 1

We equally divided 15,918 samples into two sub-cohorts. We simulated two traits using genetic effects sampled from an infinitesimal model.

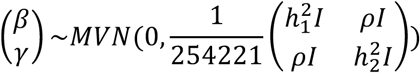

Heritability for both traits was set as 0.5. We set the genetic covariance to be 0, 0.05, 0.1, 0.15, 0.2, and 0.25.

#### Setting 2

Instead of fixing the heritability, we only assumed the heritability for both traits to be equal. Genetic correlation was fixed as 0.2. We set the genetic covariance to be 0.05, 0.1, 0.15, and 0.2, and chose heritability value accordingly.

#### Setting 3

We simulated two traits on the same sub-cohort of 7,959 samples. Heritability was fixed as 0.5 for both traits. We set the genetic covariance to be 0, 0.05, 0.1, 0.15, 0.2, and 0.25. Sample overlap correction was applied to estimate genetic covariance.

#### Setting 4

We randomly partitioned the genome into two annotation categories of the same size. We set the heritability for both traits to be 0.5, and the heritability structure does not depend on functional annotations. Genetic covariance in the first annotation was set to be 0, 0.05, 0.1, 0.15, and 0.2. Genetic effects for two traits are not correlated in the second annotation category.

#### Setting 5

We randomly partitioned the genome into three categories of the same size. Define annotation-1 to be the union of the first and the second categories, and let annotation-2 be the union of the second and the third categories. We set the heritability for both traits to be 0.5, and the heritability structure does not depend on functional annotations. Genetic covariance parameter for annotation-1 (i.e. *ρ*_1_) is set to be 0.1. We set *ρ*_2_ to be −0.2, −0.1, 0, and 0.1. The genetic covariance in regions where two annotations overlap follows an additive structure. For example, when *ρ*_1_ = 0.1 and *ρ*_2_ = −0.2, the total covariance in annotation-1 is

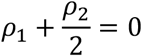

Similarly, the total covariance in annotation-2 is

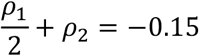

### GWAS data analysis

Details of 48 GWASs and the URLs for summary statistics files are summarized in **Table S1**. For each summary statistics dataset, we applied the same quality control steps described in [7] using the munge_sumstats.py script in LDSC. In addition, we removed all the strand-ambiguous SNPs from each dataset. For each pair of complex traits, we took the overlapped SNPs between two summary statistics files, matched the effect alleles, and removed SNPs with MAF below 5% in the 1000 Genomes Project phase-III samples with European ancestry. SNPs on sex chromosomes were also removed from the analysis. We then applied the GNOVA framework to the remaining SNPs to estimate genetic covariance. Sample overlap correction was applied when two GWASs have a large sample overlap. When calculating genetic correlation between ALS and other traits, we used previously reported 0.085 as the heritability of ALS due to negative heritability estimates [19].

### Annotation data

GenoCanyon and GenoSkyline functional annotations, as previously reported [14, 20, 21], integrate various types of transcriptomic and epigenomic data from ENCODE [22] and Roadmap Epigenomics Project [23] to predict functional DNA regions in the human genome. GenoCanyon utilizes an unsupervised learning framework to identify non-tissue-specific functional regions. GenoSkyline and GenoSkyline-Plus further extended this framework to identify tissue and cell type-specific functionality in the human genome. We applied GenoSkyline-Plus annotations for seven broadly defined tissue categories (i.e. brain, cardiovascular, epithelium, gastrointestinal, immune, muscle and other) to stratify genetic covariance by tissue type. When integrating these annotations in GNOVA, we also included the whole genome as an annotation category to guarantee that the union of all annotations covers the genome. The whole genome was not added as an additional annotation track in analyses or simulations when the functional annotations covered all SNPs in the dataset. The MAF quartiles were calculated using the genotype data of phase-III samples with European ancestry from the 1000 Genomes Project after filtering SNPs with MAF below 5%.

### LD score regression implementation

We implemented cross-trait LD score regression using the LDSC software package. For the purpose of fair comparison, we ran LD score regression on all SNPs in the dataset in the simulation studies. When analyzing real GWAS data, we followed the protocol suggested in [7] and used HAPMAP3 SNPs. LD scores were estimated using phase-I samples with European ancestry in the 1000 Genomes Project.

## Results

### Simulations

We simulated two traits using genotype data from the Wellcome Trust Case Control Consortium (WTCCC) while assuming a correlated genetic covariance structure. Detailed simulation settings are described in the **Material and methods** section. Since LDSC cannot estimate annotation-stratified genetic covariance, we compared GNOVA and LDSC using data simulated from a non-stratified, infinitesimal genetic covariance structure (**Figures 1A-D**). Both methods provided unbiased covariance estimates, but GNOVA estimator had consistently lower variance across all simulation settings. The same pattern could be observed for genetic correlation estimates (**Figure S1**). Neither method showed inflated type-I error when the true covariance is 0. When comparing the frequencies of rejecting the null hypothesis, GNOVA is nearly twice as powerful as LDSC when the true genetic covariance is below 0.1. To evaluate GNOVA’s robustness against sample overlap, we simulated two traits using genotype data of the same cohort. After applying sample overlap correction, GNOVA still outperformed LDSC, showing higher estimation accuracy and statistical power (**Figure S2**).

**Figure 1.**
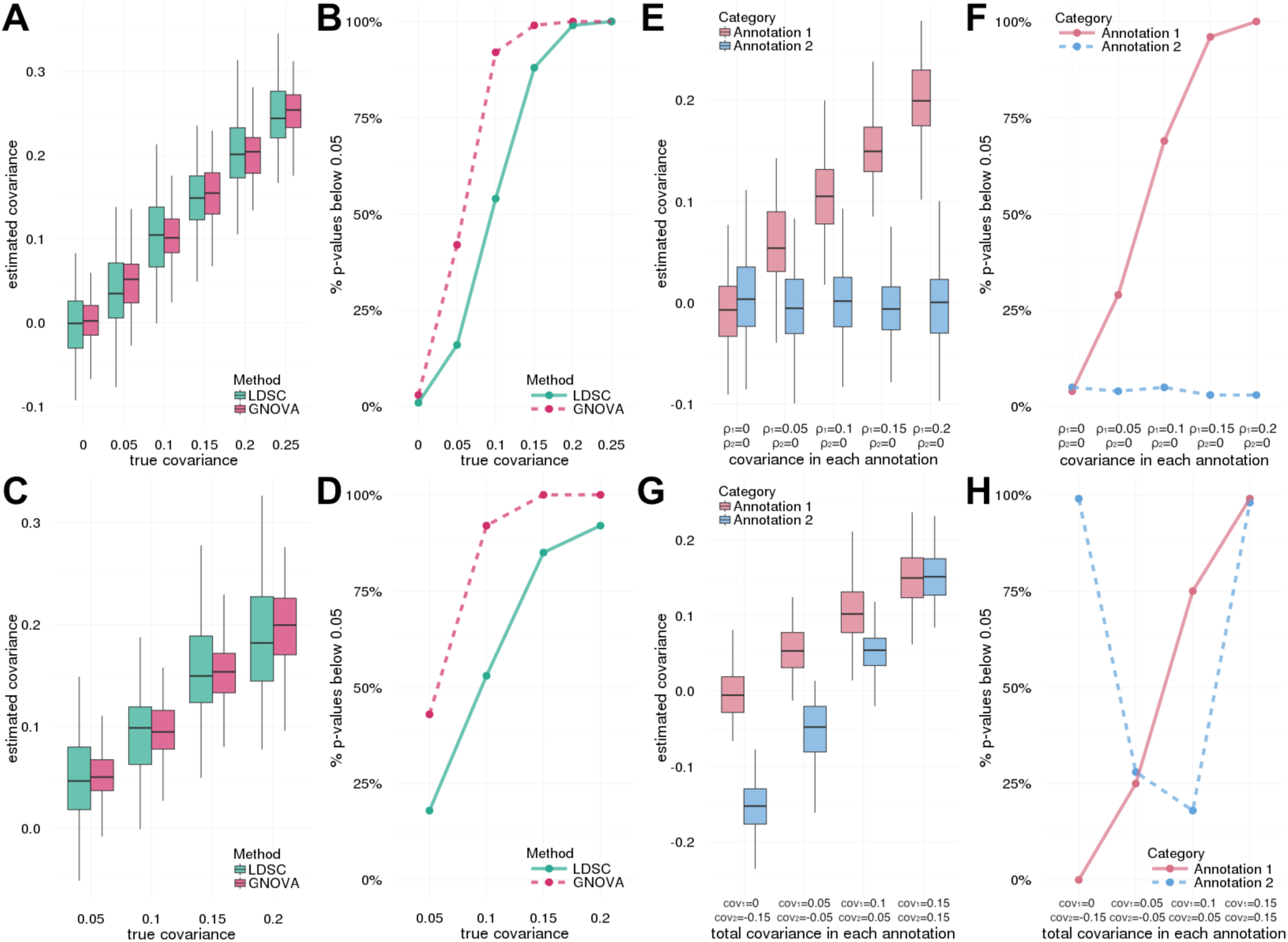
Evaluation of covariance estimation and statistical power through simulations. Detailed simulation settings are described in the **Material and methods** section. **(A-D)** Compare GNOVA and LDSC using traits simulated from a non-stratified covariance structure. We first fixed heritability for both traits but set genetic correlation to different values. The covariance estimates are shown in panel A. Panel B shows the statistical power. Next, we fixed genetic correlation but chose different values for heritability and covariance. Covariance estimates and statistical power are shown in panels C and D, respectively. **(E-H)** Estimate annotation-stratified genetic covariance. In panels E and F, we simulated data using two non-overlapping functional annotations. Results in panels G and H are based on two overlapping annotations. The true covariance values are labeled under each setting. Type-I error was not inflated when the true covariance was zero.

Next, we investigated GNOVA’s capability to estimate annotation-stratified genetic covariance. We randomly partitioned the genome into two non-overlapping annotation categories, and simulated two traits using annotation-dependent genetic covariance (**Material and methods**). GNOVA provided unbiased estimates for the genetic covariance in each category across all settings (**Figures 1E-F**). Of note, type-I error was well controlled in the annotation category without genetic covariance even when the true covariance in the other annotation category was non-zero, suggesting GNOVA’s robustness under the influence of LD. Furthermore, when functional annotations overlapped, our method still provided accurate covariance estimates and powerful inference (**Figures 1G-H**).

### Estimation of pair-wise genetic correlation for 48 human complex traits

We applied GNOVA to estimate genetic correlations for 48 complex traits using publicly available GWAS summary statistics (N_total_ ≈ 4.5 million). Trait acronyms and other details of all GWASs are summarized in **Table S1**. Out of 1,128 pairs of traits in total, we identified 176 pairs with statistically significant genetic correlation after Bonferroni correction (**Table S2** and **Figure S3**). We also applied LDSC to the same datasets and only identified 127 significant pairs (**Table S3** and **Figure S4**). A total of 52 significantly correlated trait pairs were uniquely identified by GNOVA while only 3 trait pairs were uniquely identified using LDSC. Overall, the genetic correlations estimated using GNOVA and LDSC are concordant (**Figure 2**). Consistent with our simulation results, GNOVA is more powerful when genetic correlation is moderate.

**Figure 2.**
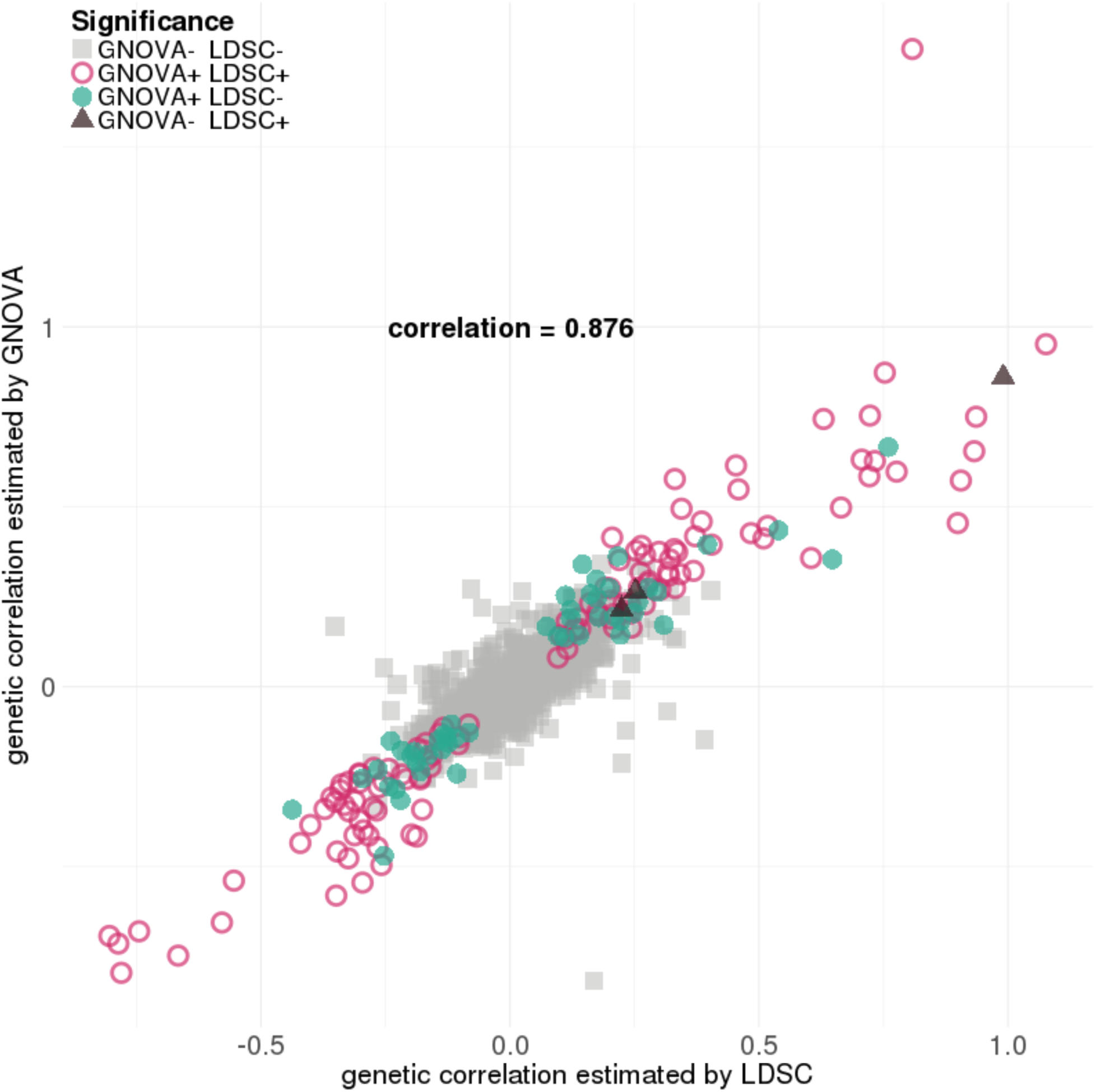
Comparison of genetic correlations estimated using GNOVA and LDSC. Each point represents a pair of traits. Overall, genetic correlation estimates are concordant between GNOVA and LDSC, but GNOVA is more powerful when genetic correlation is moderate. Color and shape of each data point represent the significance status given by GNOVA and LDSC. Trait pairs that involve gout were removed from this figure because LDSC estimated its heritability to be negative and could not properly output p-values.

To evaluate model validity, we examined correlations between several traits that are closely related either physiologically or epidemiologically (**Table S4**). As expected, femoral and lumbar bone mineral density (FNBMD and LSBMD), and depressive symptoms (DEP) and major depressive disorder (MDD [MIM: 608516]) showed strong positive genetic correlations. We also observed negative correlations between subjective well-being (SWB) and neuropsychiatric disorders such as schizophrenia [MIM: 181500], anxiety [MIM: 607834], two depression traits (DEP and MDD) and neuroticism.

We further examined pairwise correlations between 48 traits (**Figure 3**; **Figure S3**). Following hierarchical clustering, broad patterns suggesting disease relatedness emerged. These results are well documented in the literature; neuropsychiatric, metabolic diseases, and gastrointestinal inflammatory disorders clustered together with positive correlations within each individual cluster. We replicated several previous genetic correlation findings [7], including significant correlations of adult height (HGT) with coronary artery disease (CAD [MIM: 608320]) and age at menarche (AM), and of years of education (EDU) with CAD, bipolar disorder (BIP), body-mass index (BMI), triglycerides, and smoking status (SMK). Furthermore, two previous results that only passed multiple correction testing at 1% FDR passed Bonferroni correction in our analysis; namely, we observed a statistically significant negative correlation between AM and CAD, and a positive correlation between autism (ASD [MIM: 209850]) and EDU.

**Figure 3.**
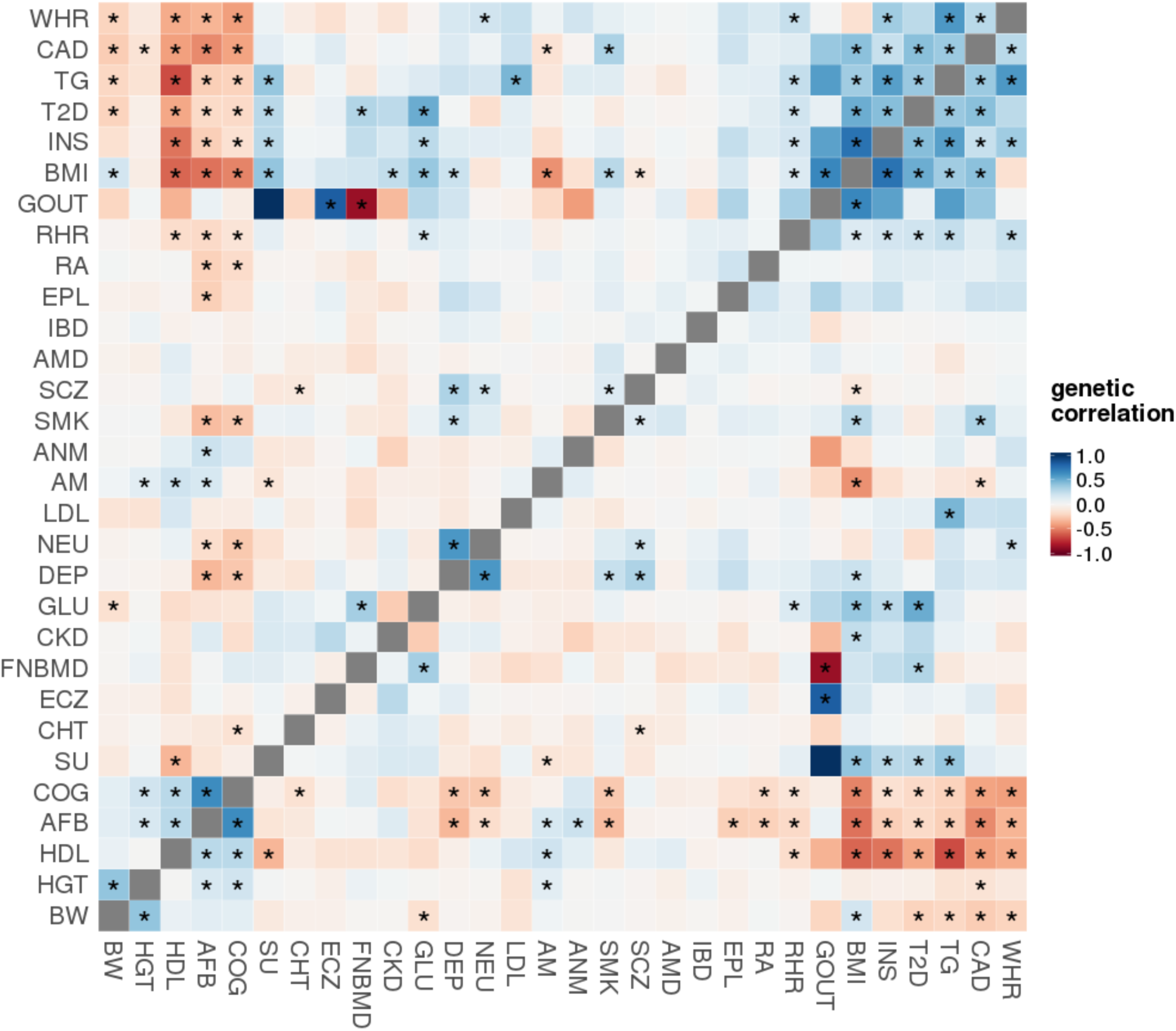
Estimated genetic correlations of 435 pairs of traits from 30 GWASs. To visualize a large number of pair-wise correlations more efficiently, we excluded closely related traits and studies with smaller sample sizes (N < 30,000) in this figure. Asterisks highlight significant genetic correlations after Bonferroni correction for all 1,128 pairs (p < 4.4×10^-5^). The complete heat map matrix is presented in **Figure S3**. The order of traits was determined by hierarchical clustering.

We also identified a number of genetic correlations that are consistent with the genetic relationships reported in the previous literature. For example, previous genetic correlation analyses identified a negative correlation between anorexia nervosa (AN [MIM: 606788]) and obesity, a result we also observed [7]. In addition, we found negative correlations of AN with glucose and triglyceride levels, as well as a positive correlation with high-density lipoprotein (HDL). These results provide further support for existing hypotheses proposing an underlying neural, rather than metabolic, etiology for metabolic syndrome [12, 21, 24]. We see an unsurprising positive correlation between glucose and insulin levels, which is consistent with our understanding of diabetes [25]. Positive correlations between multiple sclerosis (MS [MIM: 126200]) and Crohn’s disease (CD) and more generally, inflammatory bowel disease (IBD [MIM: 266600]), agree with existing reports of shared susceptibility for these diseases [26-28]. We demonstrate a positive correlation between asthma [MIM: 600807] and eczema [MIM: 603165], which share numerous loci identified in previous GWAS [29]. We also reproduced recent findings linking bone mineral density with metabolic dysfunction with positive correlations between FNBMD and both glucose and type II diabetes (T2D [MIM: 125853]) [30]. Interestingly, however, we did not see significant correlations of bone mineral density with cardiovascular diseases. Among neuropsychiatric disorders, we identified positive correlations between BIP and both depression and neuroticism. Associations between neuroticism and depression are well documented. Neuroticism is highly comorbid with MDD [31, 32], and our findings are consistent with previously observed genetic pleiotropy among neuroticism, MDD, BIP, and schizophrenia [33, 34].

Especially notable are findings that suggest a genetic basis for associations between traits regarding which the literature is either equivocal or absent, and which provide useful information to guide further study. For example, we observed correlations of serum urate (SU) with AM (-0.12), T2D (0.275), and triglycerides (0.38), and we consistently observed associations of SU and markers of metabolic syndrome. In the literature, the genetic architecture of this association has not been extensively studied [35]. Alleles in *IRF8* [MIM: 601565], a regulatory factor of type-I interferons, are associated with MS and systemic lupus erythematosus (SLE [MIM: 152700]), but with opposite effect; high type-I IFN titers are thought to be causal in SLE, but are lower in MS relative to healthy controls [36]. In this analysis, however, we found a positive correlation between MS and SLE. We also draw attention to the significant negative correlation between MS and ASD. This replicates a previous genetic association between MS and ASD, with more recent evidence suggesting shared biomedical markers, such as increase in concentrations of tumor necrosis factor-alpha (TNF-alpha), in serum in ASD and in cerebrospinal fluid in MS [37, 38]. However, previous treatment of MS with anti-TNF-alpha led to an increase in the number of demyelinating lesions and a significantly higher relapse rate [39]. Furthermore, we observed a positive genetic correlation between ulcerative colitis (UC) and primary billary cirrhosis (PBC [MIM: 109720]). CD, also an IBD and thus closely related, has been reported to share susceptibility genes with PBC including *TNFSF15* [MIM: 604052], *ICOSLG* [MIM: 605717], and *CXCR5* [MIM: 601613] [40]. Here we show that ulcerative colitis may also be genetically related to PBC.

### Stratification of genetic covariance by functional annotation

In this section, we apply functional annotations to further dissect the shared genetic architecture of 48 complex traits. We have previously developed GenoCanyon, a statistical framework to predict functional DNA elements in the human genome through integration of annotation data [20]. We partitioned the genome into two non-overlapping categories (i.e. functional and non-functional) based on GenoCanyon scores (**Material and methods**), and estimated genetic covariance within the functional and the non-functional genome for each pair of traits (**Table S5**). The total genetic covariance estimated using the stratified model is highly concordant with covariance estimated using the non-stratified model (**Figure 4A**). However, genetic covariance is enriched in the predicted functional genome for most traits (**Figure 4B**). Based on this approach, we identified one more pair of correlated traits, i.e. low-density lipoprotein (LDL) and total cholesterol (TC), whose genetic covariance largely concentrated in the predicted functional genome and achieved significance (*ρ*_*func*_ = 0.060; p = 1.0×10^-6^) while the overall covariance did not (*ρ*_*overall*_ = 0.062; p = 7.7×10^-5^).

**Figure 4.**
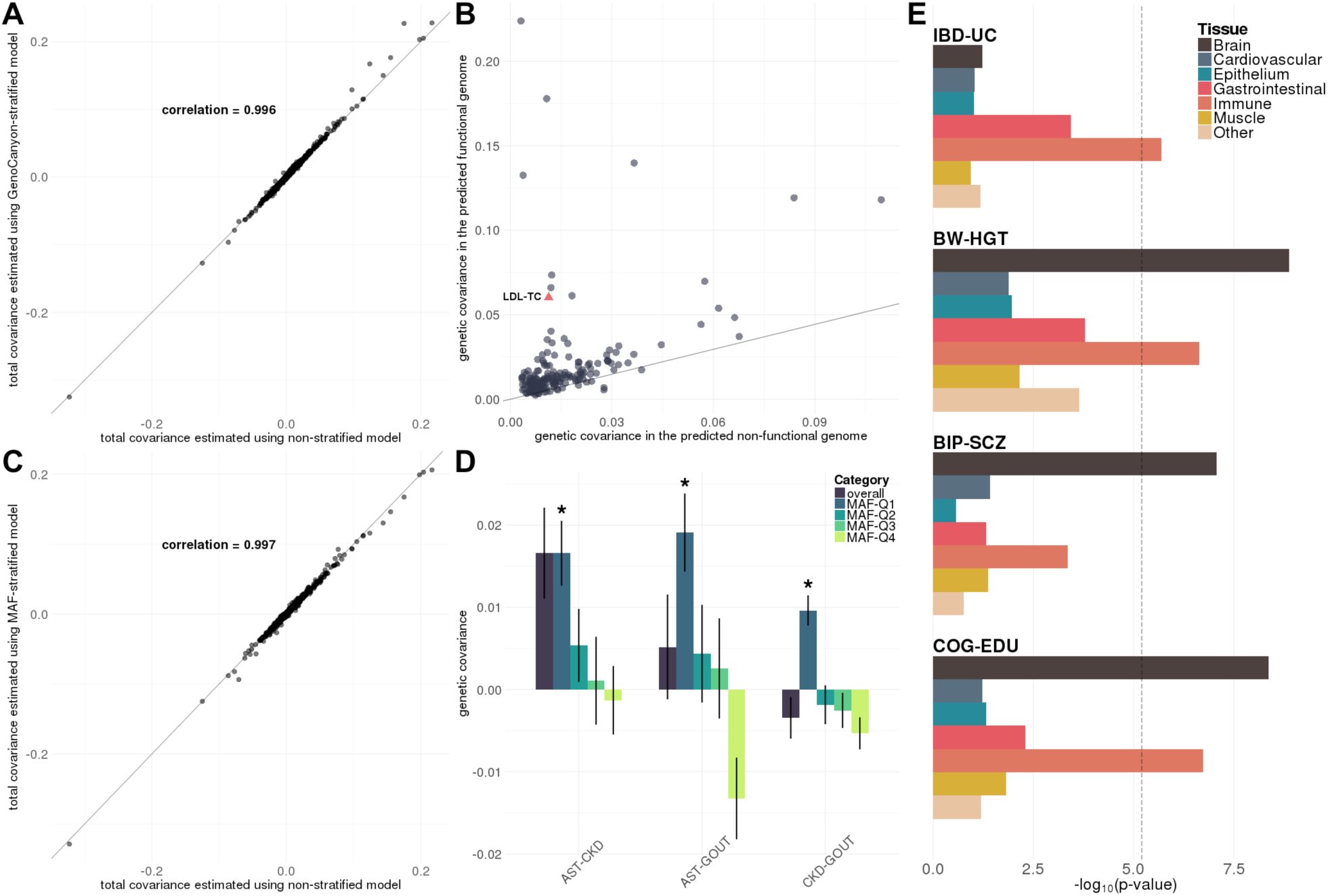
Annotation-stratified covariance analysis. **(A)** Stratify genetic covariance by genome functionality predicted by GenoCanyon. Total genetic covariance estimates were highly concordant between stratified and non-stratified models. **(B)** For significantly correlated pairs of traits based on the non-stratified model, we compared genetic covariance in the functional and the non-functional genome. Solid line marks the expected value based on annotation’s size. Trait pair LDL-TC is also plotted. **(C)** Stratify genetic covariance by MAF quartile. We compared the genetic covariance estimated by MAF-stratified and non-stratified models. **(D)** Six pairs of traits that are uniquely correlated in the lowest MAF quartile. Intervals show the standard error of covariance estimates. Asterisks indicate p-values below 4.4×10^-5^. **(E)** Stratify genetic covariance by tissue type. Each bar denotes the log-transformed p-value. Dashed line highlights the Bonferroni-corrected significance level 0.05/(7×1128) = 6.3×10^-6^.

Next, we partitioned genetic covariance based on quartiles of SNPs’ minor allele frequencies (MAFs) in subjects with European ancestry from the 1000 Genomes Project (**Material and methods**; **Table S6**). Similar to the previous analysis, we identified high concordance between the total covariance estimated using MAF-stratified model and the covariance estimates based on non-stratified model (**Figure 4C**). Overall, the estimated genetic covariance in four MAF quartiles was comparable (**Figure S5**). However, we identified three pairs of traits that are uniquely correlated in the lowest MAF quartile (**Figure 4D**), namely asthma with chronic kidney disease (CKD; p = 1.8×10^-5^), gout [MIM: 138900] with CKD (p = 4.2×10^-8^), and asthma with gout (p = 4.4×10^-5^). For several trait pairs, covariance in the lowest MAF quartile showed reversed direction compared to other quartiles. Covariance between CKD and gout even showed reversed direction compared to the estimated total covariance, highlighting the distinction in how common and less common variants are involved in the shared genetic architecture between these traits. Our findings also hint at the possible selection pressure on DNA variations contributing to metabolic traits including CKD and gout, as well as immune diseases including asthma.

Finally, we studied tissue-specificity of genetic covariance through integration of GenoSkyline-Plus annotations (**Material and methods**). GenoSkyline-Plus integrates multiple epigenomic and transcriptomic annotations from the Roadmap Epigenomics Project to identify tissue and cell type-specific functional regions in the human genome [14]. We utilized seven broadly defined tissue and cell types (i.e. brain, cardiovascular, epithelium, gastrointestinal, immune, muscle, and other) to stratify genetic covariance for 1,128 pairs of traits (**Table S7**). Six tests from 4 pairs of traits passed Bonferroni correction, i.e. p < 0.05/(1128×7) = 6.3×10^-6^ (**Figure 4E** and **Figure S6**). As expected, UC, as an IBD, was significantly and positively correlated with IBD in immune-related functional genome (p = 2.0×10^-6^); two psychiatric diseases, BIP and schizophrenia, were specifically correlated in the genome predicted to be functional in brain (p = 8.7×10^-8^). In addition, we identified cognitive function (COG) and EDU, and birth weight (BW) and HGT to be significantly correlated in both brain and immune-related functional genome. Of note, since the sizes of functional annotations are linked to statistical power, p-values here should not be interpreted as reflecting the importance of each tissue. Some tissues may be critically involved in the etiology of analyzed traits even if they may have p-values that are not statistically significant. For example, IBD and UC were substantially correlated in the gastrointestinal tract (p = 3.7×10^-4^). Many of these tests may become significant in the near future as GWASs with larger sample sizes are published.

### Dissection of shared and distinct genetic architecture between LOAD and ALS

LOAD and ALS are neurodegenerative diseases. Despite success of large-scale GWASs [19, 41], our understanding of their genetic architecture is still far from complete. We applied GNOVA to dissect the genetic covariance between LOAD and ALS using publicly available GWAS summary statistics (N_LOAD_ = 54,162; N_ALS_ = 36,052; **Table S8**).

We identified positive and significant genetic correlation between LOAD and ALS (correlation = 0.175, p = 2.0×10^-4^). LDSC provided similar estimates but failed to achieve significance (**Table 1**). 82.6% of the total genetic covariance between LOAD and ALS is concentrated in 33% of the genome predicted to be functional by GenoCanyon (p = 8.2×10^-5^). Furthermore, MAF-stratified analysis showed that 54.6% of the covariance could be explained by the SNPs in the highest MAF quartile (p = 0.005). In fact, genetic covariance is lower with lower MAF, and covariance in the lowest MAF quartile is nearly negligible. This is surprising considering that the heritability of ALS is enriched in variants with lower MAF [19]. We also performed tissue-stratified analysis using GenoSkyline-Plus annotations (**Table S9**). No tissue passed the significance threshold after multiple testing correction, but covariance is more concentrated in immune, brain, and cardiovascular functional genome, and showed nominal significance in the immune annotation track (p = 0.014). Whether this will lead to a potential neuroinflammation pathway shared between LOAD and ALS remains to be studied in the future using larger datasets.

**Table 1.** Dissection of genetic covariance between LOAD and ALS. Numbers in parentheses indicate standard errors. Significant p-values after adjusting for multiple testing within each section are highlighted in boldface.

| Annotation | Category | Covariance | P-value |
| --- | --- | --- | --- |
| Non-stratified | GNOVA | 0.016 (0.004) | **$2.0 \times 10^{-4}$** |
|  | LDSC | 0.012 (0.007) | 0.075<sup>a</sup> |
| GenoCanyon | functional | 0.016 (0.004) | **$8.2 \times 10^{-5}$** |
|  | non-functional | 0.003 (0.004) | 0.377 |
| MAF | Q1 | -0.001 (0.003) | 0.842 |
|  | Q2 | 0.003 (0.004) | 0.361 |
|  | Q3 | 0.004 (0.004) | 0.327 |
|  | Q4 | 0.008 (0.003) | **0.005** |

<sup>a</sup> p-value in LDSC was calculated from genetic correlation instead of genetic covariance.

Next, we stratified genetic covariance between LOAD and ALS by chromosome. Somewhat surprisingly, we did not observe a linear relationship between per-chromosome genetic covariance and chromosome size (**Figure 5A**) given that the overall genetic covariance is positive and significant. Since we have observed the concentration of genetic covariance in the functional genome, we further partitioned each chromosome by genome functionality. We identified a clear and positive linear relationship between genetic covariance in the functional genome and the size of predicted functional DNA on each chromosome (**Figure 5B**). The correlation between per-chromosome genetic covariance in the non-functional genome and the size of non-functional chromosome is negative and significantly smaller than the corresponding quantity in the functional genome (**Figure S7**; p = 0.044; tested using Fisher transformation). Our findings suggest a polygenic covariance architecture between LOAD and ALS, and highlight the importance of stratifying genetic covariance by functional annotation.

**Figure 5.**
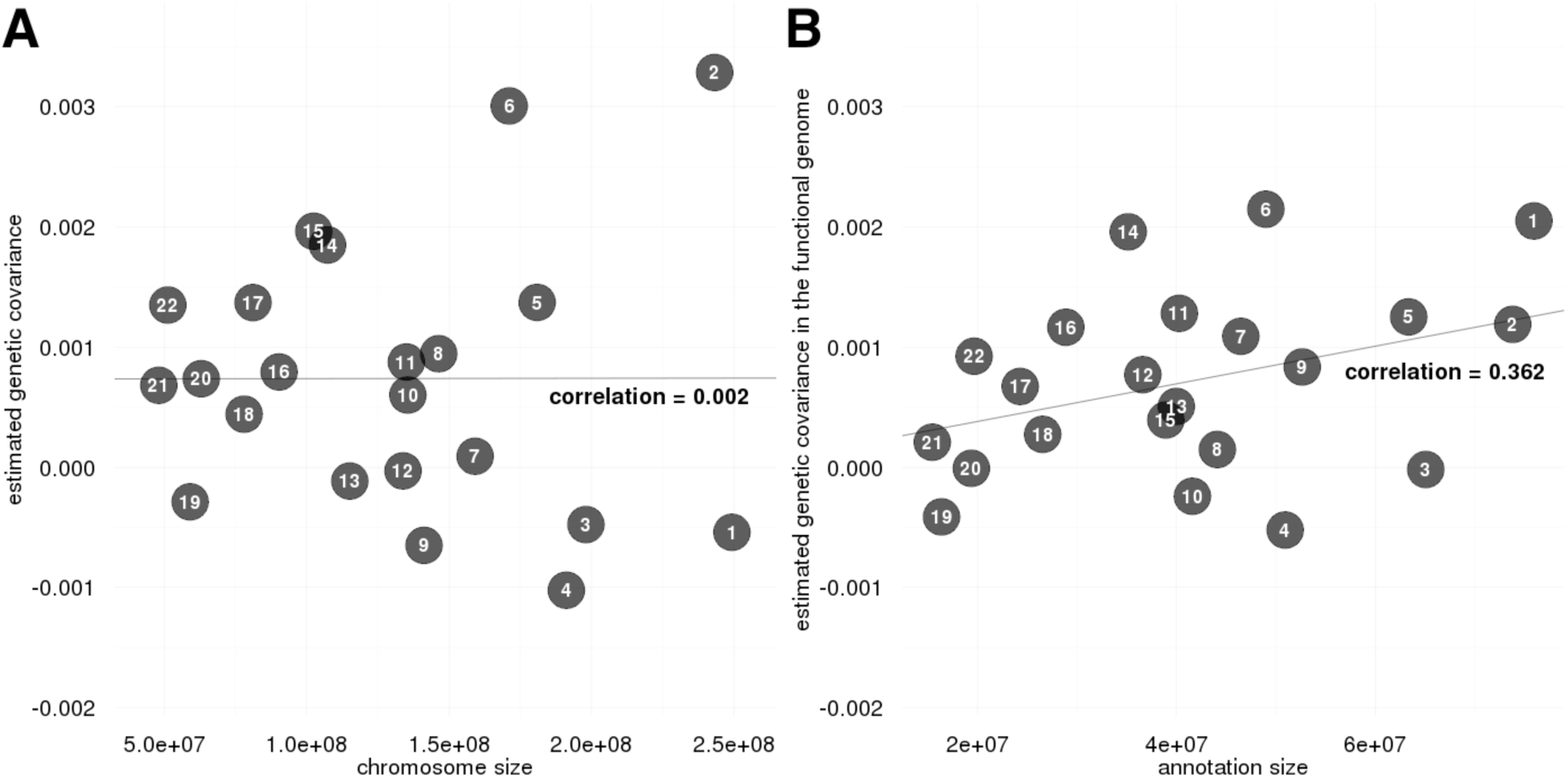
Stratification of genetic covariance between LOAD and ALS by chromosome. **(A)** Comparisons of the estimated per-chromosome genetic covariance with chromosome size. **(B)** Comparisons of the estimated genetic covariance in the predicted functional genome on each chromosome with size of the functional genome.

Finally, we jointly analyzed LOAD, ALS, and 48 other complex traits (**Table S10**). Interestingly, LOAD and ALS showed distinct patterns of genetic correlations with other complex traits (**Figure 6**). We identified negative and significant correlations between LOAD and cognitive traits including COG and EDU. HGT and age at first birth (AFB), two traits related to hormonal regulation as well as socio-economic status, were also significantly and negatively correlated with LOAD. Consistent with previous reports, we did not identify substantial correlation between LOAD and other neurological and/or psychiatric diseases [7, 9]. We identified negative correlations between LOAD and gastrointestinal inflammatory diseases including a significant correlation with PBC. Asthma and eczema were both positively correlated with LOAD, suggesting a complex genetic relationship between LOAD and different immune-related diseases. Although some of these traits had the same correlation direction with ALS, none of them was significant. Instead, ALS was significantly and positively correlated with MS, a neurological disease with a well-established immune component [42]. ALS was also positively correlated with several other immune-related diseases including celiac disease (CEL [MIM: 212750]), asthma, PBC, and IBD (including CD and UC), though none of these was statistically significant. The nominal correlations between ALS and neurological and psychiatric diseases including epilepsy, schizophrenia, BIP, AN, and MDD also remain to be validated in the future using studies with larger sample sizes.

**Figure 6.**
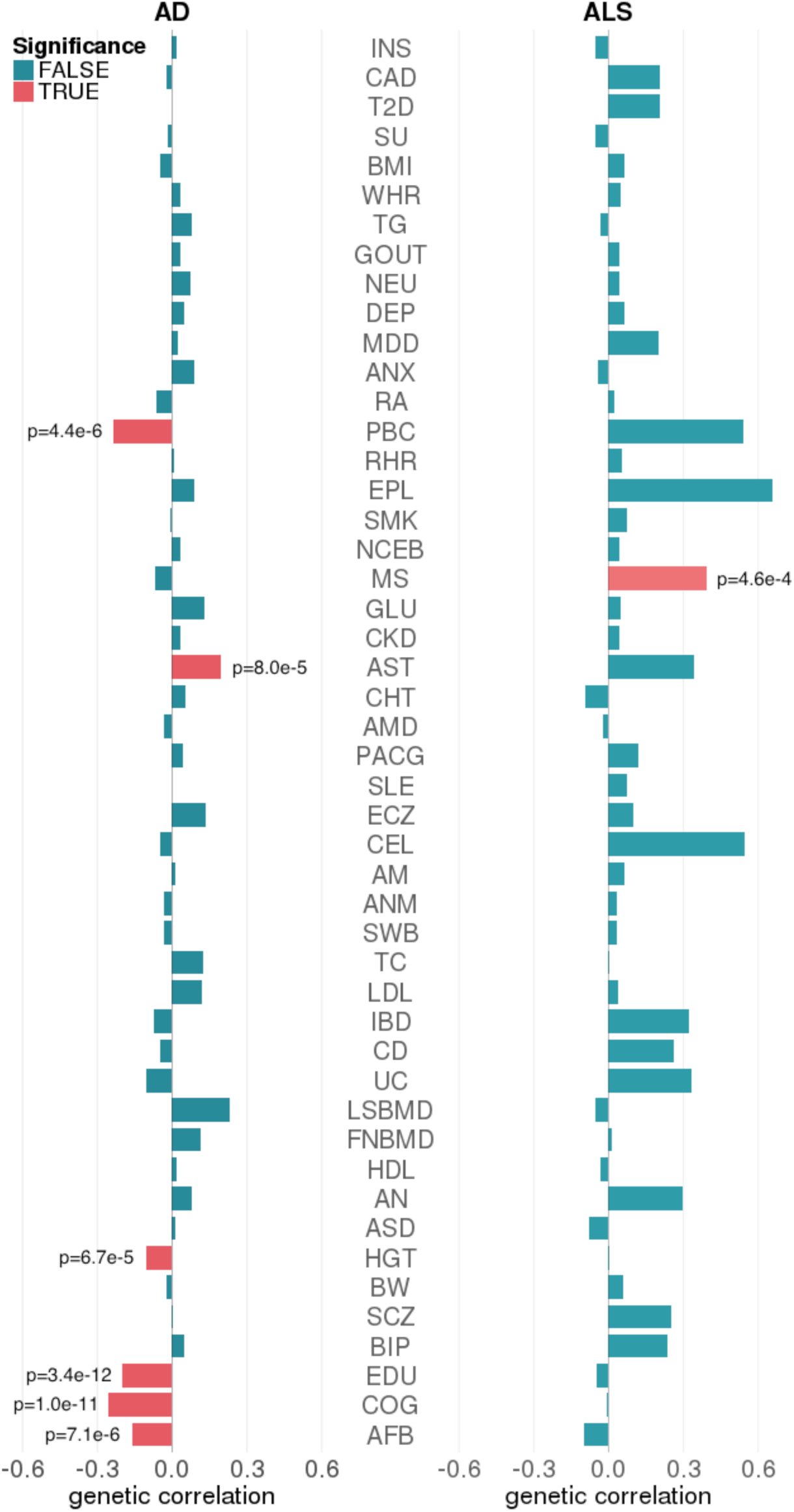
Genetic correlations between LOAD, ALS, and 48 complex traits. Significant pairs with p < 0.05/(48×2) = 5.2×10^-4^ are highlighted in red.

## Discussion

Although our understanding of complex disease etiology is still far from complete, we have gained valuable knowledge on the genetic architecture of numerous complex traits from large-scale association studies, partly due to advances in statistical genetics. First, a large proportion of trait heritability can be explained by SNPs that do not pass the Bonferroni-corrected significance threshold [1]. Therefore, it is often helpful to utilize genome-wide data instead of only focusing on significant SNPs in post-GWAS analyses. Second, sample size is critical for many statistical genetics applications. However, individual-level genotype and phenotype data from consortium-based GWASs are not always easily accessible due to policy and privacy concerns. Thanks to the great efforts from large international collaborations such as the Psychiatric Genomics Consortium in promoting open science and data sharing, it has become a tradition for GWAS consortia to share summary statistics to the broader scientific community. Therefore, it is of practical interest to use GWAS summary statistics as the input of downstream analytical methods [8]. Finally, integration of high-throughput transcriptomic and epigenomic annotation data has been shown to improve statistical power as well as interpretability in many recent complex trait studies [12-14]. As large consortia such as ENCODE [22] and Roadmap Epigenomics Project [23] continue to expand, integrative approaches based on functional genome annotations will become an even greater success. In this paper, we developed a novel method to estimate and partition genetic covariance between complex traits. Our method enjoys all the aforementioned advantages. It only requires genome-wide summary statistics and a reference panel as input, and allows stratification of genetic covariance by functional genome annotation, which provides novel insights into the shared genetic basis between complex traits and in some cases, improves the statistical power.

Numerous studies have hinted at a shared genetic basis among neurodegenerative diseases [43, 44]. Due to the convenience and efficiency of LDSC and the wide accessibility of GWAS summary statistics, several attempts have been made to estimate genetic correlation between neurodegenerative diseases [9, 45]. To date, these efforts have not been as successful as similar studies on psychiatric diseases and immune-related traits. One reason is that existing methods may not be statistically powerful enough to identify moderate genetic correlation using GWASs with limited sample sizes. In addition, the shared genetics among neurodegenerative diseases may not fit the global, infinitesimal covariance structure that most existing tools are based on. In this study, we applied GNOVA to dissect the genetic covariance between LOAD and ALS, two major neurodegenerative diseases, using summary statistics from the largest available GWASs. Our findings suggest that covariance between LOAD and ALS is concentrated in the predicted functional genome and in very common SNPs. Moreover, after applying functional annotations to stratify the genome, estimated per-chromosome genetic covariance is proportional to chromosome size, suggesting a shared polygenetic architecture between LOAD and ALS and also demonstrating the importance of incorporating predicted genetic activity with GenoCanyon. In addition, joint analysis with 50 complex traits also revealed distinctive genetic covariance profiles for LOAD and ALS. LOAD is negatively correlated with multiple traits related to cognitive function and hormonal regulation, while ALS is positively correlated with MS and a few other immune-related traits. Our findings provided novel insights into the shared and distinct genetic architecture between LOAD and ALS, and also further demonstrated the benefits of incorporating functional genome annotations into genetic covariance analysis.

Also of note are findings involving serum urate. SU was positively correlated with gout but also with a few metabolic traits. Gout is an arthritic inflammatory process caused by deposition of uric acid crystals in joints, and the role of hyperuricemia in gout is well established. More recently, a role for hyperuricemia in the pathophysiology of metabolic syndrome and CKD has been suggested [46]. While associations between hyperuricemia and cardiovascular disease are well described [47], multiple hypotheses exist regarding details of its involvement [48]. For example, hyperuricemia may lead to inflammation in the kidney through vascular smooth muscle proliferation, inducing hypertension via pre-glomerular vascular changes [49]. It has also been shown to induce oxidative stress in various settings; in adipocytes and islet cells, this may be involved in development of diabetes, and it may also result in impaired endothelin function and activation of the renin-angiotensin-aldosterone system, leading to hypertension [50-53]. Despite this evidence, genetic investigations have not identified a strong relationship between hyperuricemia and metabolic syndrome. Polymorphism in gene *SLC22A12* [MIM: 607096] was associated with hyperuricemia but not with metabolic syndrome [54]. Mendelian randomization studies showed an association between uric acid and gout, but did not find an association with T2D, or cardiovascular risk factors such as hypertension, glucose, or CAD [55, 56]. Our results suggest that GNOVA successfully isolated a signal of biological and clinical significance that provides important impetus for further inquiry in the etiology of metabolic syndrome.

Dissecting relationships among complex traits is a major goal in human genetics research. Genetic covariance is a useful metric to quantify such relationships, but it has its limitations. First, genetic covariance implicitly imposes a strong assumption on the shared genetic basis between complex traits. Not only may the same set of genetic components affect multiple traits, their effect sizes on both traits are also assumed to be proportional. In the future, it is of interest to extend our method to estimate more generalized metrics, e.g. consistency in effect directions. Second, genetic covariance analysis does not highlight specific DNA segments with pleiotropic effects. Several SNP-based methods have been developed to identify pleiotropic associations using GWAS summary statistics [57, 58]. However, due to the large number of SNPs in the genome, statistical power is a critical issue and large-scale inference remains challenging. In addition, we have demonstrated that integrating functional annotations into genetic covariance analysis could reveal subtle structures in shared genetics between complex traits, but interpretation of genetic covariance remains a challenge. Pickrell et al. recently proposed an approach to distinguishing causal relationship among traits from pleiotropic effects via independent biological pathways [59]. Han et al. developed a method to distinguish pleiotropy from phenotypic heterogeneity [60]. Although many questions remain unanswered, these recent studies have broadened our view on interpreting complex genetic relationships between human traits. Further, statistical power in genetic covariance analysis will be reduced if the shared genetic components have discordant effect directions on different traits. This problem can be partly addressed by the aforementioned SNP-based methods. Recently, Shi et al. developed a method to estimate local heritability and genetic correlation [61, 62]. This approach provides an alternative methodological option for analyzing genetic effects at specific loci. Finally, we note that common SNPs in GWAS do not fully explain phenotypic similarity. For example, the estimated genetic covariance among lipid traits only explains 10∼15% of their phenotypic covariance available on LD Hub [10]. Other factors such as rare variants, copy number variations, and environmental factors may have substantial contributions to the phenotypic covariance among complex traits. Dissection of these complex relationships will be an interesting topic to pursue in the future. Our method, in conjunction with many other tools, provides the most complete picture to date about shared genetics between complex phenotypes.

In summary, we developed GNOVA, a novel statistical framework to perform powerful, annotation-stratified genetic covariance analysis using GWAS summary statistics. Through theoretical proof, we have established GNOVA’s statistical optimality within the framework of method of moments. Compared to LD score regression, GNOVA provides more accurate genetic covariance estimates and powerful statistical inference. Its unique feature of performing annotation-stratified analysis also adds depth to existing analysis strategies. Using GNOVA, we were able to expand the discovery of genetic covariance among a spectrum of common diseases and complex traits. Our findings shed light onto the shared and distinct genetic architecture of complex traits. As the sample sizes in genetic association studies continue to grow, our method has the potential to continue identifying shared genetic components and providing novel insights into the etiology of complex diseases.

## Supplemental data description

Supplemental data include complete notes for mathematical details, ten figures, and ten tables.

## Acknowledgements

This study was supported in part by the National Institutes of Health grants R01 GM59507, the VA Cooperative Studies Program of the Department of Veterans Affairs, Office of Research and Development, and the Yale World Scholars Program sponsored by the China Scholarship Council. Dr. Crane’s and Dr. Mukherjee’s efforts were supported by grant R01 AG042437 and U01 AG006781.

This study makes use of data generated by the Wellcome Trust Case-Control Consortium. A full list of the investigators who contributed to the generation of the data is available from www.wtccc.org.uk. Funding for the project was provided by the Wellcome Trust under award 076113, 085475, and 090355. We thank the International Genomics of Alzheimer’s Project (IGAP) for providing summary results data for these analyses. The investigators within IGAP contributed to the design and implementation of IGAP and/or provided data but did not participate in analysis or writing of this report. IGAP was made possible by the generous participation of the control subjects, the patients, and their families. The i-Select chips were funded by the French National Foundation on Alzheimer’s disease and related disorders. EADI was supported by the LABEX (laboratory of excellence program investment for the future) DISTALZ grant, Inserm, Institut Pasteur de Lille, Université de Lille 2, and the Lille University Hospital. GERAD was supported by the Medical Research Council (Grant n° 503480), Alzheimer’s Research UK (Grant n° 503176), the Wellcome Trust (Grant n° 082604/2/07/Z), and German Federal Ministry of Education and Research (BMBF): Competence Network Dementia (CND) grant n° 01GI0102, 01GI0711, 01GI0420. CHARGE was partly supported by the NIH/NIA grant R01 AG033193 and the NIA AG081220 and AGES contract N01–AG–12100, the NHLBI grant R01 HL105756, the Icelandic Heart Association, and the Erasmus Medical Center and Erasmus University. ADGC was supported by the NIH/NIA grants: U01 AG032984, U24 AG021886, U01 AG016976, and the Alzheimer’s Association grant ADGC–10–196728. We are also grateful for all the consortia and investigators that provided publicly accessible GWAS summary statistics.

The authors declare no conflict of interests.

## Ethical statement

Procedures followed were in accordance with the ethical standards of the responsible committee on human experimentation. Proper informed consent was obtained when needed.

## Author contribution

Q.L. and H.Z. conceived and designed the study.

Q.L., B.L., D.C., and W.D. performed the statistical analyses.

Q.L., B.L., D.O., and Y.H. developed the statistical model and studied its theoretical properties.

M.E. assisted in interpreting genetic correlations.

R.L.P. and T.J. developed the software package.

Q.L., B.L., Y.H., D.C., W.D., Q.H., and Z.L. implemented the algorithm.

B.L. and C.J. implemented jackknife estimation.

Q.L., M.E., and H.Z. wrote the manuscript.

S.M. and P.K.C. advised on the analysis of neurodegenerative diseases.

H.Z. advised on statistical and genetic issues.

All authors read and approved the manuscript.

## Web resources

GNOVA

https://github.com/xtonyjiang/GNOVA

LDSC

https://github.com/bulik/ldsc/

LD Hub

http://ldsc.broadinstitute.org/ldhub/

OMIM

http://www.omim.org

